# Investigating Short-Chain Fatty Acids Effects on Extracellular Vesicles Production in Colorectal Cancer

**DOI:** 10.1101/2024.10.30.620636

**Authors:** D. Macedo, S. Abalde-Cela, L. Diéguez, A. Preto, C. Honrado

## Abstract

Colorectal cancer (CRC) is the third most diagnosed and the second leading cause of cancer-related deaths globally, often due to late detection and limited treatment options. Recent studies have linked alterations in gut microbiota to CRC, particularly emphasizing the role of short-chain fatty acids (SCFAs) like acetate, propionate, and butyrate in shaping the tumor microenvironment (TME). SCFAs contribute to CRC pathogenesis by inducing lysosomal membrane permeabilization, cell cycle arrest, and apoptosis in cancer cells. Extracellular vesicles (EVs) are membrane-bound vesicles that facilitate intercellular communication and have gained attention as promising non-invasive biomarkers for cancer diagnosis and treatment monitoring. EVs participate in cellular response mechanisms to external stimuli by transferring proteins, lipids, and nucleic acids between cells, thus modulating target cell behavior and promoting coordinated responses to stress and environmental challenges. This process is essential for cellular adaptation and plays a significant role in pathophysiological processes, including tumor progression and immune modulation, making EVs highly relevant in clinical research. This study examined the impact of SCFAs on EV production and phenotype in CRC cells. The results indicated a notable increase in EV-sized particles following SCFA treatment of colorectal cell lines, particularly in the SW480 CRC cell line. For CRC cell lines, while co-precipitated protein levels remained stable, there was a slight decrease in cellular DNA and an increase in EV-associated DNA. KRAS-mutant SW480 cells exhibited the most pronounced response, emphasizing their heightened sensitivity to SCFA. Notably, microsatellite instability - a key biomarker for immunotherapy in CRC - was detected in both small and large EV populations from BRAF-mutant RKO cells after SCFA treatment, even at low DNA concentrations. These findings underscore the potential of EVs for non-invasive detection of molecular markers, paving the way for further exploration of their role in precision oncology.

Graphical Abstract.
Effects of SCFA on EV production and characteristics in CRC cells. Treatment with SCFA led to a significant increase in the number of EV-sized particles, a decrease in cellular DNA and a corresponding increase in EV-DNA. This study also identified MSI in both s-EV and L-EV, even following SCFA treatment and at low DNA concentrations. Created using BioRender.

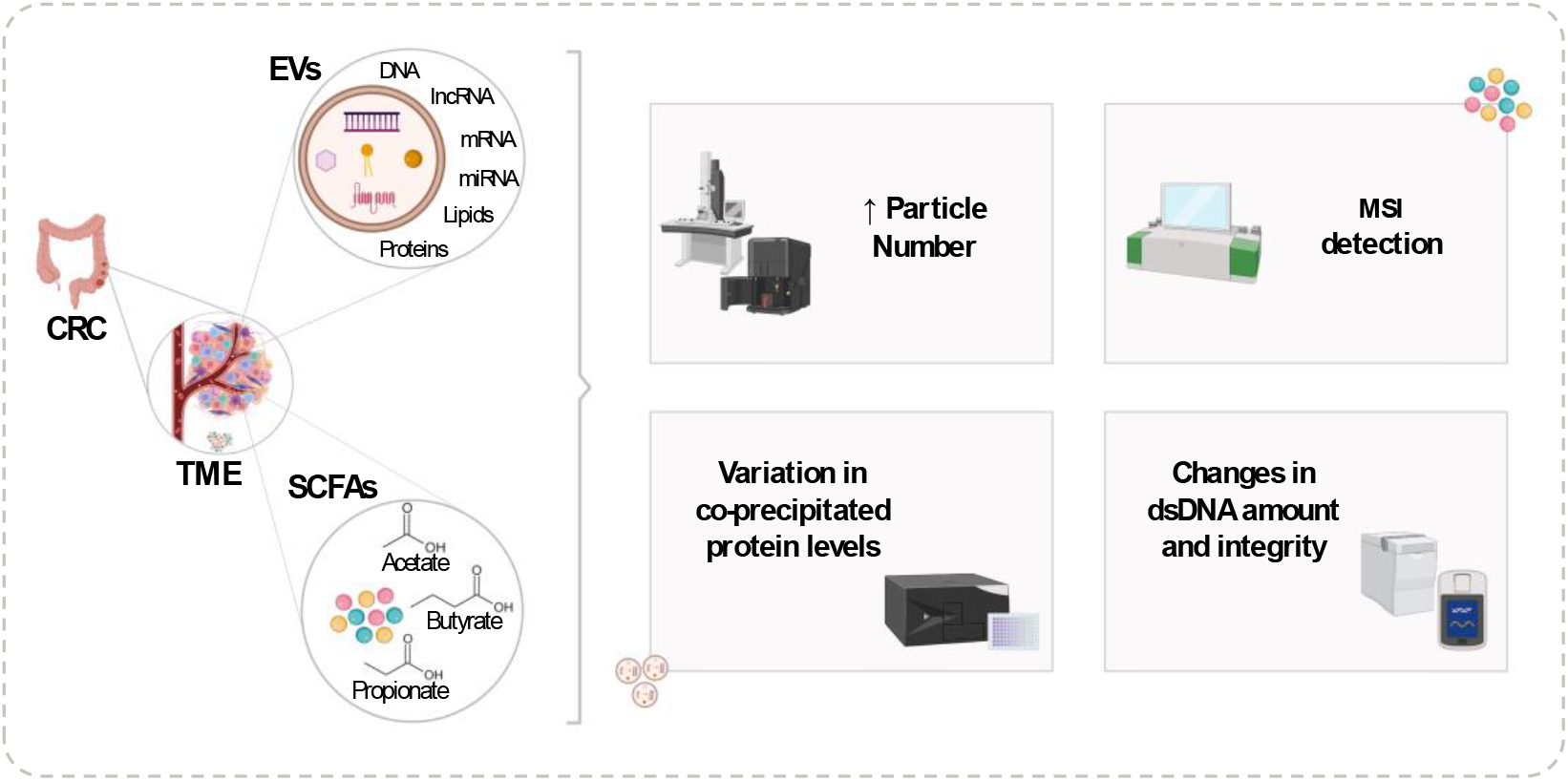

## Introduction

Colorectal cancer (CRC) remains one of the most prevalent and lethal cancers globally, ranking as the third most diagnosed and second leading cause of cancer-related deaths. The high mortality rate associated with CRC is often due to late-stage diagnosis and limited treatment options. This has intensified the search for innovative approaches in early detection, accurate prognosis, and novel therapies (1). Emerging research points to a strong link between gut microbiota alterations and CRC pathogenesis, with particular focus on the role of short-chain fatty acids (SCFAs) in modulating the tumor microenvironment (TME) (2).

SCFAs, including acetate, propionate, and butyrate, are produced by gut microbiota through the fermentation of dietary fibers. These metabolites play a key role in maintaining intestinal homeostasis and have been shown to exhibit anticancer properties in CRC through several mechanisms. For instance, butyrate, known as a histone deacetylase (HDAC) inhibitor, influences gene expression by promoting acetylation, which can induce cell cycle arrest and apoptosis in CRC cells (3). Both acetate and propionate have been shown to decrease cellular proliferation and to induce lysosomal membrane permeabilization, mitochondrial dysfunction, and oxidative stress, thus promoting cancer cell death. Additionally, SCFAs exhibit anti-inflammatory effects by modulating cytokine production and immune cell infiltration in the TME, suggesting their role in reducing pro-tumorigenic inflammation (4,5). Despite these promising effects, the exact mechanisms by which SCFAs influence tumor progression and their potential applications in CRC management remain areas of active research.

In parallel with these findings, extracellular vesicles (EVs) have earned considerable attention in the context of cancer biology. EVs are small (mostly < 1 µm), membrane-bound phospholipid vesicles released by cells into the extracellular space, carrying a diverse range of biomolecules such as proteins, lipids, and nucleic acids. These vesicles play a crucial role in intercellular communication and have been shown to reflect the physiological and pathological states of their parent cells. Their ability to circulate in bodily fluids, coupled with their potential to carry cancer-specific markers, positions EVs as promising candidates for non-invasive cancer diagnostics, including liquid biopsy approaches (6,7). In particular, the role of EVs in CRC is increasingly recognized, as they have been shown to contribute to tumor growth, metastasis, and immune evasion (8). Moreover, tumor-derived EVs may carry markers from their cells of origin with valuable clinical importance, such as DNA carrying, for example, KRAS mutation, which can inform therapy decisions, namely regarding anti-EGFR therapies, which are ineffective in patients harboring these mutations (9,10). Additionally, EV-DNA may contain microsatellite instabilities (MSI), which have been shown to be an effective predictive biomarker for immune checkpoint inhibitors, making MSI evaluation a good candidate for immunotherapy selection (11,12). Still, the clinical application of EVs is currently limited by a lack of standardized methods for purifying and analyzing EVs, hindering the translation of EV-based diagnostics into clinical practice (7). Despite these challenges, EVs remain a promising avenue for improving CRC detection and management.

Given the emerging importance of SCFAs in CRC therapy and the growing recognition of EVs as clinical tools, this study seeks to explore how SCFAs influence the production and molecular profile of EVs released by CRC cells. Investigating the interactions between SCFAs and EVs could offer new insights into the development of non-invasive clinical tools and potential therapeutic strategies in CRC.

## Materials and Methods

### 1. Cell Culture Conditions

This study used three cell lines: SW480 and RKO, derived from human colorectal cancer, and NCM460, a normal colon mucosal epithelial cell line. SW480 cells harbor KRAS^G12V^ and TP53^R273H/P3095^ mutations, while RKO cells possess BRAF^V600E^ and PIK3A^H1047R^ mutations. SW480 and RKO cell lines were obtained from IPATIMUP (Porto, Portugal), and NCM460 cell line was obtained from INCELL (San Antonio, Texas, USA). SW480 and NCM460 cells were cultured in RPMI 1640 medium with stable Glutamine, while RKO cells were grown in DMEM High Glucose supplemented with stable Glutamine and Sodium Pyruvate. Both media were supplemented with 10 % fetal bovine serum (FBS) and 1 % penicillin/streptomycin. Cells were maintained at 37 °C in a humidified incubator with 5 % CO_2_. Conditioned media (CM) was produced by seeding cells in 150 mm dishes, allowing them to reach 50 %–70 % confluence after 48 hours, then washing, adding fresh medium, and incubating for an additional 48 hours before collection.

### 2. Short-chain fatty acids treatment

A solution of short-chain fatty acids was prepared using sodium acetate (CAT #S2889), sodium propionate (CAT #P1880), and sodium butyrate (CAT #303410), all sourced from Sigma-Aldrich (Burlington, Massachusetts, USA). The SCFAs were combined in a molar ratio of 60:25:15 (acetate:propionate:butyrate), reflecting the physiological concentrations found in the colon under normobiosis. Stock solutions of SCFAs were prepared weekly, diluted in sterile deionized water, and filtered for sterility.

The half-maximal inhibitory concentrations (IC50), as well as the IC25 and IC10 values for the SCFA mixture, were previously established by our research group (**Table 1**). Conditioned media for EV isolation was generated by seeding the cells as described earlier and adding the IC50 concentrations of SCFAs to the medium at 48 hours. After an additional 48 hours, CM was collected, and cell counts were assessed.

**Table 1.** IC10, IC25 and IC50 concentrations (mM) of SCFA mix (3)

|  | NCM460 | RKO | SW480 |
| --- | --- | --- | --- |
| IC<sub>10</sub> | 5.05 | 16.92 | 3.19 |
| IC<sub>25</sub> | 13.39 | 28.51 | 5.36 |
| IC<sub>50</sub> | 36.65 | 47.61 | 7.30 |

### 3. EV Isolation

After collecting CM, a standard centrifugation protocol was performed, involving centrifugation at 500×g for 10 minutes to remove dead cells, and at 2,000×g for 10 minutes to eliminate cell debris, with collection of the supernatant after each step. This was followed by centrifugation at 10,000×g for 20 minutes at 4 °C, where the supernatant was collected for small EVs isolation, and the pellet was retained as the fraction containing large EVs. Subsequently, the supernatant went to 2 h of Ultracentrifugation at 120,000×g, followed by the removal of the supernatant until 1.5 mL remained, addition of 18.5 mL of PBS 1× and another ultracentrifugation at 120,000 × *g* for 2 hours (“2 × UC 2 h”). After the UC steps, the supernatant was removed, and 1 mL of the pellet was washed with 500 µL of PBS 1 ×, collected, and frozen.

### 4. Particle morphology, number and size

To analyze the morphology and size of EVs from all cell lines, TEM was employed due to its high resolution, allowing for the visualization of nanoscale structures. Each EV sample was prepared from 160 mL of CM, which was subjected to UC for 16 hours at 110,000×g. The supernatant was discarded, and 18.5 mL of filtered PBS was added to the 1.5 mL pellet. Following this, a second UC was performed, at 110,000×g for 6 hours. The resulting pellet (12 mL) was then subjected to UF to concentrate the EVs down to a final volume of 750 µL. Of this concentrated sample, 500 µL was used for SEC, where the first 4 fractions were collected, yielding a final volume of 1.6 mL. For imaging, Quantifoil holey Carbon grids were used (Quantifoil, Jena, Germany). A sample of 5 µL from the undiluted EV sample was allowed to settle on the carbon surface of the grid for 2 minutes. The samples were then stained twice with 2 % uranyl acetate for 30 seconds to enhance contrast. The prepared grids were imaged using a JEOL JEM-2100 electron microscope (JEOL Ltd., Akishima, Tokyo, Japan), equipped with a LaB6 electron gun operating at 200 kV. A magnification range of 40,000 × to 80,000 × was employed to allow for detailed observation of the EVs.

EV size, size distribution and concentration were evaluated using Nanoparticle Tracking Analysis (NTA) on the NanoSight NS300 (Malvern, Worcestershire, United Kingdom) equipped with a low-volume flow cell. Samples were diluted in filtered dPBS to a final volume of 1 mL, targeting a particle concentration of 15 to 45 particles per frame. Optimal collection and analysis settings were optimized with a Gain of 1, Camera Level of 14, and Detection Threshold of 14. Each dilution was measured at least three times for 60 seconds using a continuous flow pump set at 40.

The size and size distribution of large EVs samples, obtained post-centrifugation at 10,000×g, were analyzed using Dynamic Light Scattering (DLS) on a Litesizer 500 (Anton Paar, Graz, Austria) equipped with a 648 nm, 40 mW laser. The samples were diluted 5-fold in filtered dPBS to reach a final volume of 1 mL, achieving a transmittance between 70 % and 80 % for all measurements. This ensured an appropriate particle concentration, allowing for accurate size and distribution measurements.

### 5. Co-precipitated protein levels

Protein quantification was carried out using the Pierce BCA Protein Assay Kit (Thermo Fisher Scientific, Massachusetts, USA), which detects and quantifies protein levels based on the reduction of Cu^2+^ to Cu^1+^ in the presence of peptides in an alkaline medium. The cuprous cations (Cu^1+)^ are then detected colorimetrically using a reagent containing bicinchoninic acid. Standards and working reagents were prepared following the manufacturer’s instructions. Briefly, 25 µL of each standard or sample was pipetted in triplicate into a microplate well, followed by the addition of 200 µL of Working Solution to each well. The plate was covered, mixed on a plate shaker, and incubated at 37 °C for 30 minutes. Lastly, absorbance was read at 562 nm using the BioTeK Synergy H1 (Winooski, Vermont, USA) microplate reader.

### 6. dsDNA extraction and quantification

DNA was isolated from the EV samples using the QIAamp DNA Mini Kit (Qiagen, Hilden, Germany), in accordance with the manufacturer’s instructions. To summarize, 20 µL of Proteinase K and 200 µL of Buffer AL were combined to 200 µL of each sample, followed by vortexing for 15 seconds. The samples were then incubated at 56 °C for 10 minutes, after which 200 µL of ethanol (96 %) was added. The resulting mixture was transferred to a spin column and centrifuged at 6,000×g for 1 minute. The flow-through was discarded and 500 µL of Buffer AW1 was added, followed by another round of centrifugation at 6,000×g for 1 minute, with the flow-through again discarded. Subsequently, 500 µL of Buffer AW2 was added, and the sample was centrifuged at 20,000×g for 3 minutes. The spin column was transferred to a new 2 mL collection tube, and 200 µL of Buffer AE was added. After centrifugation at 6,000×g for 1 minute, the flow-through, containing the DNA, was collected. In some cases, deoxyribonuclease (DNase I, Molecular Biology, Thermo Fisher Scientific, Massachusetts, USA) was included prior to the extraction to remove cell-free DNA.

The extracted DNA was quantified using the Qubit Fluorometer 4.0 and the 1× dsDNA High Sensitivity Quantification Kit (both from Invitrogen, California, USA), following the manufacturer’s instructions. Briefly, standards were prepared and read, and 1-20 µL of each sample was then added to 180-199 µL of the Working Solution, making a final volume of 200 µL. The samples were left at room temperature for 2 minutes before their fluorescence was measured using the equipment.

### 7. Integrity of dsDNA

The quality of the extracted DNA was evaluated using the Agilent 2100 Bioanalyzer system, with the Agilent DNA 12000 Kit (both from Agilent, Technologies, California, USA), according to the manufacturer’s instructions. In short, the gel-dye mix was prepared by combining 25 µL of the DNA dye concentrate to a DNA gel matrix vial, vortexing the mixture, transferring it to a spin filter and centrifuging at 1,500×g for 10 minutes. Afterwards, the Gel-Dye Mix was loaded, as well as the marker, ladder, and the samples, corresponding to DNA previously extracted and quantified, as described in Section 7.3. The chip was then evaluated in the 2100 BioAnalyzer.

### 8. Presence of Microsatellite Instability

Samples from NCM460, RKO, and SW480 were analyzed with the Bio-Rad ddPCR MSI Detection Kit (Bio-Rad Laboratories, California, USA), including cells, large EVs, and small EVs, under control conditions and after treatment with SCFA, with and without DNase application. The kit was used according to the manufacturer’s instructions. For each assay, two controls were required: a No Template Control (NTC) and a Positive Control (PC). Reaction mixtures were prepared in labeled tubes, combining nuclease free-water, ddPCR Multiplex Supermix, and Assay (1, 2, or 3). Each mixture was aliquoted into designated wells, with 6.6 µL of NTC, PC, and sample DNA added as required. Unused wells were filled with water and ddPCR Buffer Control. The plate was sealed with a foil heat seal and then heat sealed at 180 °C for 5 seconds. The corners of the plate were vortexed and the plate was centrifuged for 1 minute at 1,150×g. Droplet generation was performed using the Automated Droplet Generator according to operational manuals. The plate was then transferred to a thermal cycler for PCR amplification using the specified cycling conditions. After completion, the plate was held at 4 °C until the lid temperature cooled to 37 ºC and could be stored at 2 °C to 8 °C overnight before analysis on the QX600 Droplet Reader (Bio-Rad Laboratories, California, USA).

### 9. Data Analysis

All statistical data presented were obtained from a minimum of three independent experiments and are expressed as mean ± standard deviation (SD). The statistical analyses were conducted employing either unpaired t tests, one-way ANOVA and Dunnett’s or Tukey’s multiple comparisons tests, or two-way ANOVA with Sidak’s or Tukey’s multiple comparisons tests. The analysis was carried out using the GraphPad Prism Software (version 8.4.3 for Windows, GraphPad Software, San Diego, CA, USA). The significance of values was established according to the following p values: * p ≤ 0.05; ** p ≤ 0.01; *** p ≤ 0.001; **** p ≤ 0.0001 for a confidence level of 95 %.

For the BioAnalyzer results, a script was developed in VisualStudioCode using Python to normalize the Excel data, convert it from seconds to base pairs, and establish the base pairs peak and fragmentation ranges, ultimately plotting the resulting size distribution graph.

## Results and Discussion

### 1. Particle morphology, concentration and size distribution

The small EV (s-EV) particle counts measured by Nanoparticle Tracking Analysis (NTA) revealed a marked increase in EV production across all cell lines following SCFA treatment (**Figure 1 A**). Specifically, NCM460, RKO, and SW480 cell lines exhibited 1.86-, 1.38-, and 2.15-fold increases, respectively, potentially reflecting SCFA-induced apoptotic activity. This increase in EV output may enhance the potential for molecular characterization, facilitating the identification of disease-associated biomarkers or therapeutic responses.

**Figure 1.**
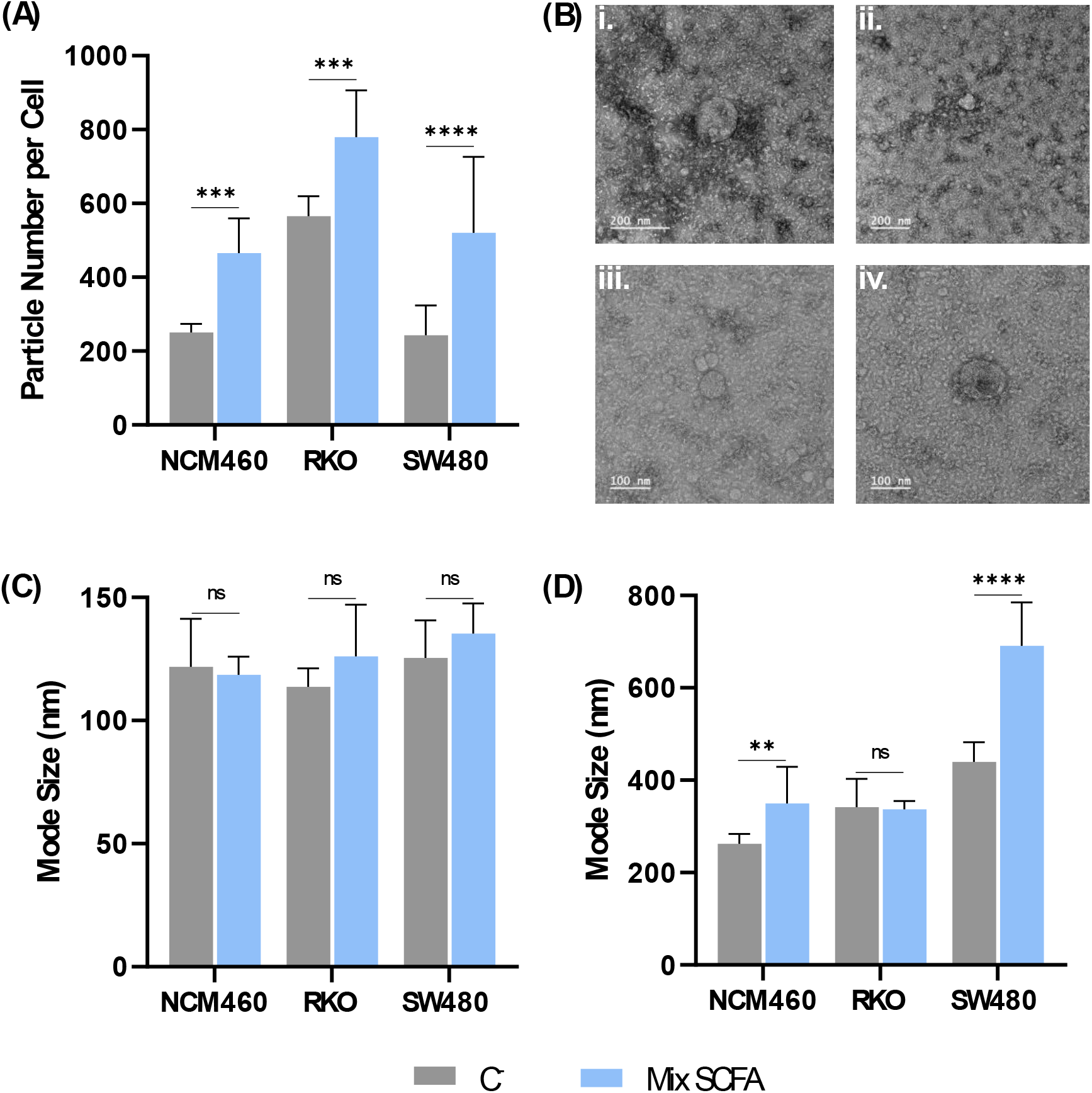
Impact of SCFA treatment on EV production and size. For each cell line, (A) quantification of s-EV particle counts per cell; (B) representative TEM images of s-EVs from NCM460 (i and ii), RKO (iii), and SW480 cells (iv); and mode size of (C) s-EVs and (D) L-EVs, measured by NTA and DLS, respectively. Results are presented as mean ± SD for each measurement, from at least three independent experiments. Statistical analysis was conducted using two-way ANOVA and Sidak’s multiple comparisons tests. ** p< 0.01; *** p<0.001; **** p< 0.0001.

Transmission electron microscopy (TEM) images (**Figure 1 B**) displayed s-EV particle sizes consistent with the literature (13), and NTA measurements confirmed the mode sizes (**Figure 1 C**) aligned with TEM results. There were no significant size differences between EVs derived from SCFA-treated and untreated cells. However, tumor-derived EVs from SCFA-treated cells were slightly larger than those from the non-tumorigenic NCM460 cell line. This slight increase may correlate with apoptotic EV release, as apoptotic events are known to produce larger vesicles containing cellular components reflective of stress responses.

Quantification of large EVs (L-EVs) was limited by the upper detection threshold of NTA (1 µm) (14). Therefore, dynamic light scattering (DLS) was used to assess L-EV size (**Figure 1 D**). Although DLS does not yield accurate concentration data, due to its reliance on intensity-weighted averages skewed by larger particles (14), it confirmed an increase in L-EV size after SCFA treatment in NCM460 and SW480 cell lines. This effect could result from SCFA-induced stress responses, such as histone deacetylase (HDAC) inhibition, which promotes apoptosis and autophagy, leading to membrane remodeling and the release of larger vesicles, including apoptotic bodies (15). In contrast, no significant size changes were detected in EVs from the RKO cell line, possibly due to distinct molecular characteristics, mutation profiles, and different responses to SCFA-induced stress (16).

### 2. Co-precipitated protein levels

The anticancer effects of SCFAs in CRC cells are associated with mechanisms like cell cycle arrest, lysosomal membrane permeabilization, cytosolic acidification, and induction of apoptosis (2). Consistent with these effects, no significant differences were observed in extracellular protein levels in the CRC cell lines SW480 and RKO following SCFA treatment (**Figure 2**). This likely indicates that cellular components are encapsulated within apoptotic bodies, thus minimally impacting extracellular protein concentrations. In contrast, a reduction in protein concentration was detected in the large EV fraction from the epithelial cell line NCM460 after SCFA treatment, which may be attributed to SCFA-induced autophagy. In normal cells, SCFA can promote autophagy as part of their anti-inflammatory effects, leading to increased intracellular degradation and recycling rather than extracellular release of cellular proteins (17,18). In the s-EV fraction, a slight increase in co-precipitated protein content was noted (p=0.0176), potentially due to partial EV disruption during isolation. Additionally, despite thorough purification with two rounds of ultracentrifugation (2 × UC 2 h), some extracellular proteins may have remained in the conditioned media.

**Figure 2.**
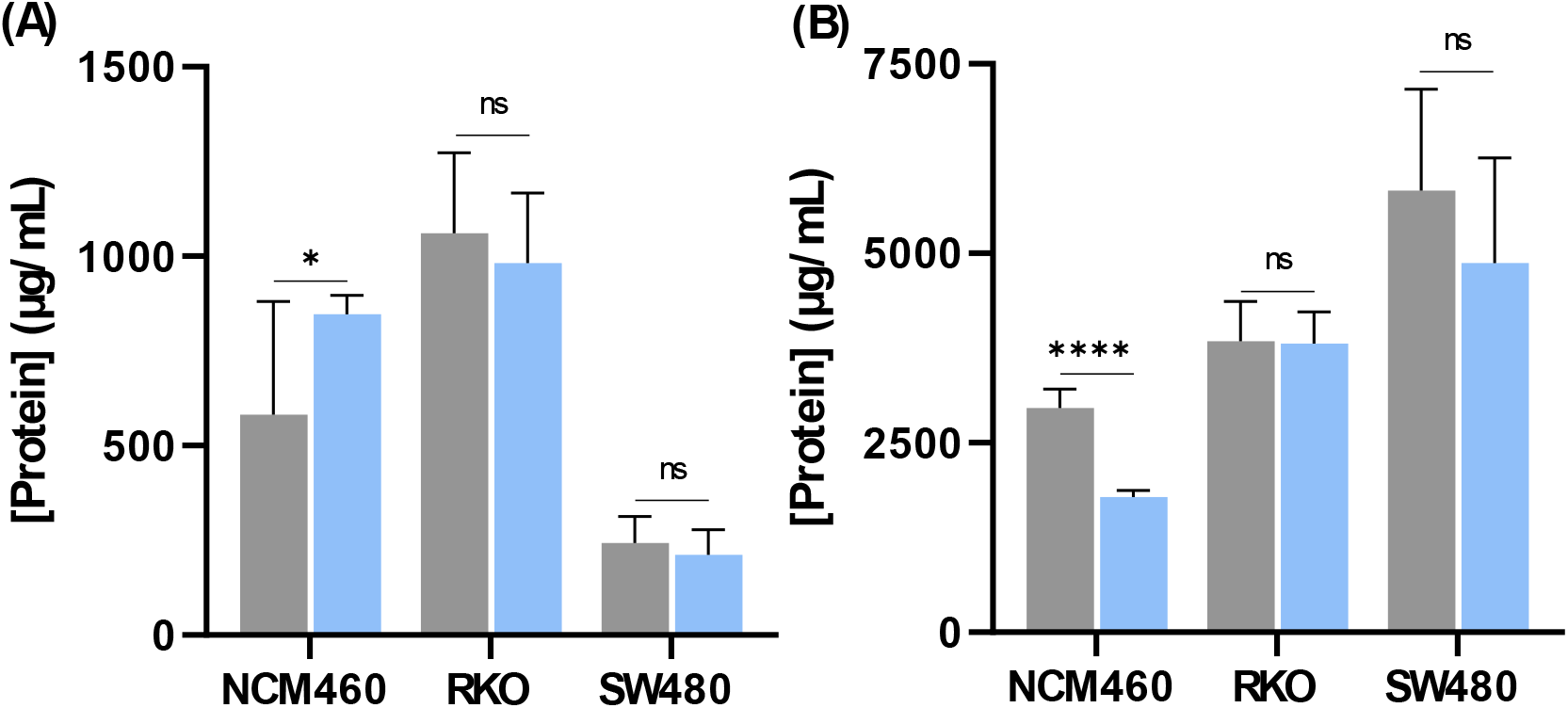
Effect of SCFAs on the concentration of co-precipitated proteins in EV samples. Co-precipitated protein levels were quantified by BCA for both (A) s-EVs and (B) L-EVs from each cell line under conditions with no treatment and with SCFA treatment. Results are expressed as mean ± SD for each measurement, from at least three independent experiments. Statistical analysis was conducted using two-way ANOVA and Sidak’s multiple comparisons tests. * p<0.05; ** p< 0.01; *** p<0.001; **** p< 0.0001.

### 3. Quantification of dsDNA

EVs are key in biomedical research and clinical applications due to their biocompatibility and ability to carry internal cargoes, including DNA. Compared to cell-free DNA or circulating tumor DNA, EV-derived DNA (EV-DNA) remains more stable due to the protective effect of the EV lipid membrane, which shields the internal cargoes from removal agents in their surrounding environment. As a result, EV-DNA closely reflects the genetic profile of tumor cells. This increases the potential for reliable, non-invasive access to tumor-specific genetic material, making EVs promising candidates for biomarker discovery (19,20). Accordingly, this study quantified and analyzed EV-DNA following SCFA treatment to investigate SCFA-induced effects on cellular mechanisms and evaluate EV-DNA integrity. DNA was quantified in cells (**Figure 3 A**), large EVs (**Figure 3 B**), and small EVs (**Figure 3 C**) from the NCM460, RKO, and SW480 cell lines, under both control and SCFA-treated conditions, with and without DNase I treatment to eliminate free DNA and isolate EV-DNA.

**Figure 3.**
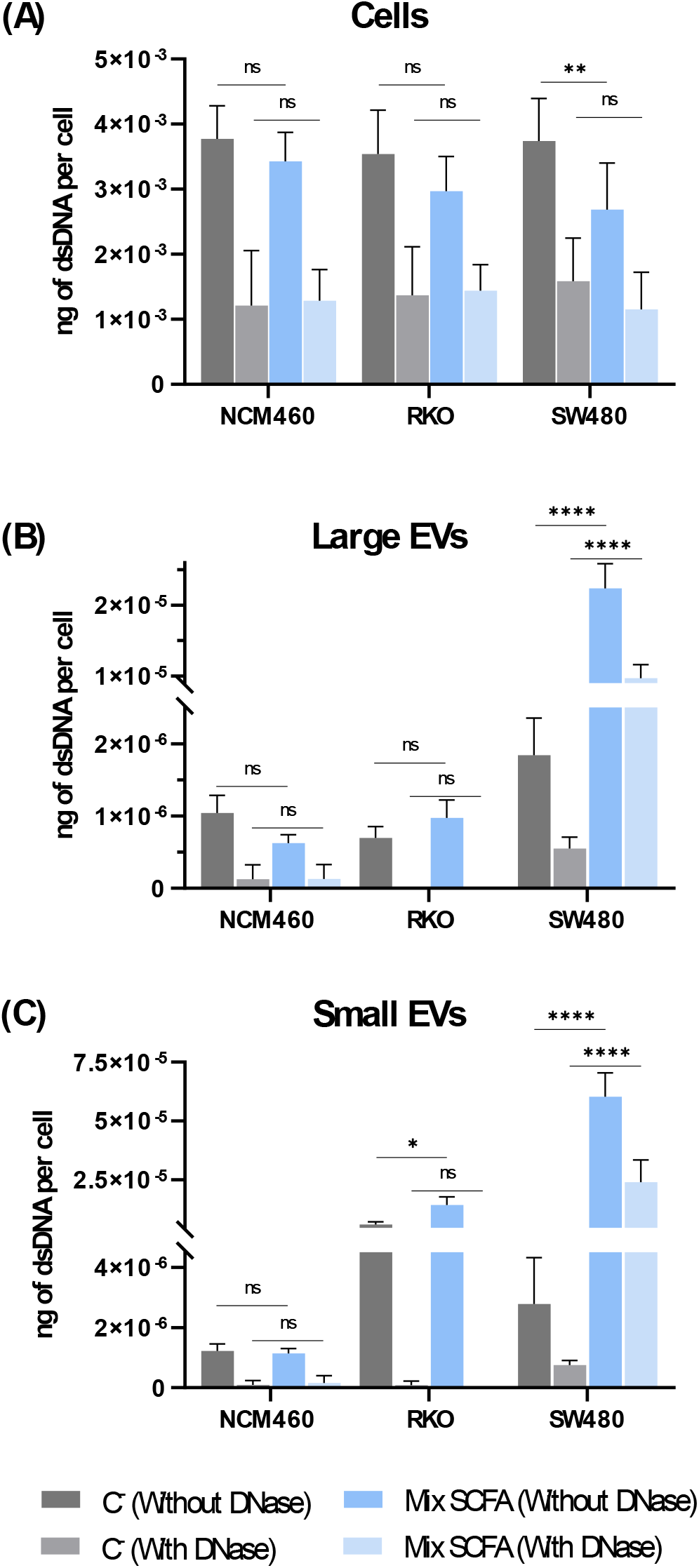
Effect of SCFAs on dsDNA quantification. The quantification of dsDNA was quantified with and without DNase treatment, under both the control and SCFA treatment conditions, across all cell lines for their corresponding (A) cells, and (B) large and (C) small EVs. Measurements were conducted using Qubit Fluorometer 4.0, with the 1× HS dsDNA Quantification Kit. Results are expressed as mean ± SD for each measurement, from at least three independent experiments. Statistical analysis was performed using two-way ANOVA and Tukey’s multiple comparisons tests. * p<0.05; ** p< 0.01; *** p<0.001; **** p< 0.0001.

Following DNase treatment, a significant reduction in dsDNA levels was observed across all samples, indicating the presence of free DNA in initial measurements. Following SCFA treatment, and with DNase application, cellular dsDNA levels remained mostly unchanged (**Figure 3 A**). In contrast, without DNase, a slight decrease in cellular DNA was observed, most noticeably in cancer cell lines RKO and SW480 (p-values of 0.2099 and 0.0026, respectively), suggesting that DNA is being packaged into apoptotic bodies rather than released extracellularly as cell-free DNA (21).

Regarding EV-DNA quantification (**Figure 3 B** and **3 C**), only EVs from the SW480 cell line contained sufficient dsDNA for measurement after DNase treatment. In SW480 cells, SCFA treatment led to a 17.7-fold increase in large EV-DNA and a 32.8-fold increase in small EV-DNA, confirming that SCFAs stimulate the production of EVs carrying DNA, likely through apoptotic body formation. In the absence of DNase, slight increases in dsDNA were observed in both RKO and SW480 EVs, with the SW480 line showing a more substantial rise. These differences may be due to differences in the genetic and metabolic profiles of the cell lines (22). Meanwhile, NCM460 cells exhibited no significant change in small EV-DNA and a slight reduction in large EV-DNA, aligning with SCFAs’ possible autophagic effects in normal epithelial cells, which promote intracellular recycling rather than DNA release (16,17).

These results (**Figure 3**), combined with particle counts per cell (**Figure 1 A**) and co-precipitated protein concentration measurements (**Figure 2**), highlight a complex relationship between SCFA-induced cellular responses and EV characteristics. Notably, SW480 cells, with their lower IC50 values (**Table 1**), demonstrated heightened sensitivity to SCFA treatment, reflected by significant increases in EV counts and EV-DNA content. This suggests SCFAs induce apoptotic pathways in SW480 cells, leading to EVs being enriched with genetic material. In contrast, RKO cells showed more moderate increases in EV numbers and dsDNA, indicating a comparatively lower SCFA sensitivity, while NCM460 cells exhibited no notable EV-DNA changes, possibly due to SCFA-induced autophagy. These findings collectively suggest that SCFA-induced stress responses in CRC cells result in differential EV release and composition, with DNA-containing EVs potentially serving as novel diagnostic or prognostic markers.

### 4. dsDNA Integrity

Assessing DNA integrity is critical for evaluating genomic stability, cellular damage, and therapeutic efficacy (23). As previously mentioned, SCFAs exert anticancer effects by inducing apoptosis via HDACs inhibition, which promotes chromatin relaxation and DNA fragmentation (2). DNA integrity analysis, particularly within EVs, could thus provide key insights into cancer cell states and treatment impacts (24). In this study, DNA integrity was analyzed using Bioanalyzer, which employs capillary electrophoresis for precise DNA fragment size assessment based on migration speeds. While effective, the Bioanalyzer’s sensitivity limit of 0.5 ng/µL posed challenges for EV-derived samples with low DNA concentrations (25).

To analyze fragmentation patterns, a custom Python script was developed for electrophoresis data, enabling the alignment of fluorescence units and quantification of fragment regions relative to a specific peak. While SCFA treatment induced slight DNA fragmentation in cell samples (observed through a shift toward smaller fragments and a decrease in base-pair peak), no bands were detectable in EV-DNA samples, likely due to low DNA quantities (**Appendix 1**). This trend aligns with existing data and suggests that SCFA exposure may fragment DNA in CRC cells, but the RKO cell line showed resistance, possibly due to reduced sensitivity to SCFAs or diminished fluorescence intensity, which complicated analysis.

### 5. Presence of Microsatellite Instability

MSI is a key biomarker in CRC, indicating genetic instability due to DNA mismatch repair deficiencies. MSI-high (MSI-H) tumors typically have a positive prognosis and lower metastasis rates, but often show poor responses to conventional chemotherapy (26). Hence, MSI testing is essential for making treatment decisions, particularly for immunotherapy. Traditional MSI detection methods include PCR and immunohistochemistry, standards in clinical practice. Recent advances in liquid biopsy technology enabled MSI detection in ctDNA and in cancer cells, providing a non-invasive way to monitor tumor dynamics (27). Exploring MSI within EVs also represents a promising area of research, that could yield valuable insights into tumor behavior. While ctDNA-based MSI detection is feasible, its high degree of fragmentation reduces the likelihood of identifying specific fragments containing MSI biomarkers. Additionally, traditional biopsy-based DNA analysis is invasive and complex to implement in clinical practice. In contrast, analyzing MSI within EVs offers a non-invasive alternative that simplifies sample collection, preserves genetic material, and enhances the accurate detection of disease biomarkers, with potential for improved patient stratification and management.

Droplet digital PCR (ddPCR) was selected for MSI detection in EV-DNA due to its sensitivity and precision in detecting low-frequency variants in cfDNA, which is particularly useful in liquid biopsies. The Bio-Rad ddPCR kit for MSI detection uses five specific markers: BAT25 and BAT26 (Assay 1), NR21 and NR24 (Assay 2), and Mono27 (Assay 3). In clinical settings, utilizing this pentaplex panel, a tumor is classified as MSI if at least three out of five markers are positive (11).

Following DNA extraction and quantification, ddPCR was performed on DNA from cells, L-EVs, and s-EVs, under control and SCFA-treated conditions, both with and without DNase. Samples were validated by a minimum of 10,000 droplets to ensure reliable detection. The MSI status of each sample was determined using the relative abundance of mutant versus wild-type DNA copies. Consistent with earlier reports (22), ddPCR confirmed MSI in RKO cells and microsatellite stability (MSS) in SW480, while providing, for the first time, a confirmation on the MSS status of the non-cancerous NCM460 cell line, as well as in their respective large EVs (**Figure 4**).

**Figure 4.**
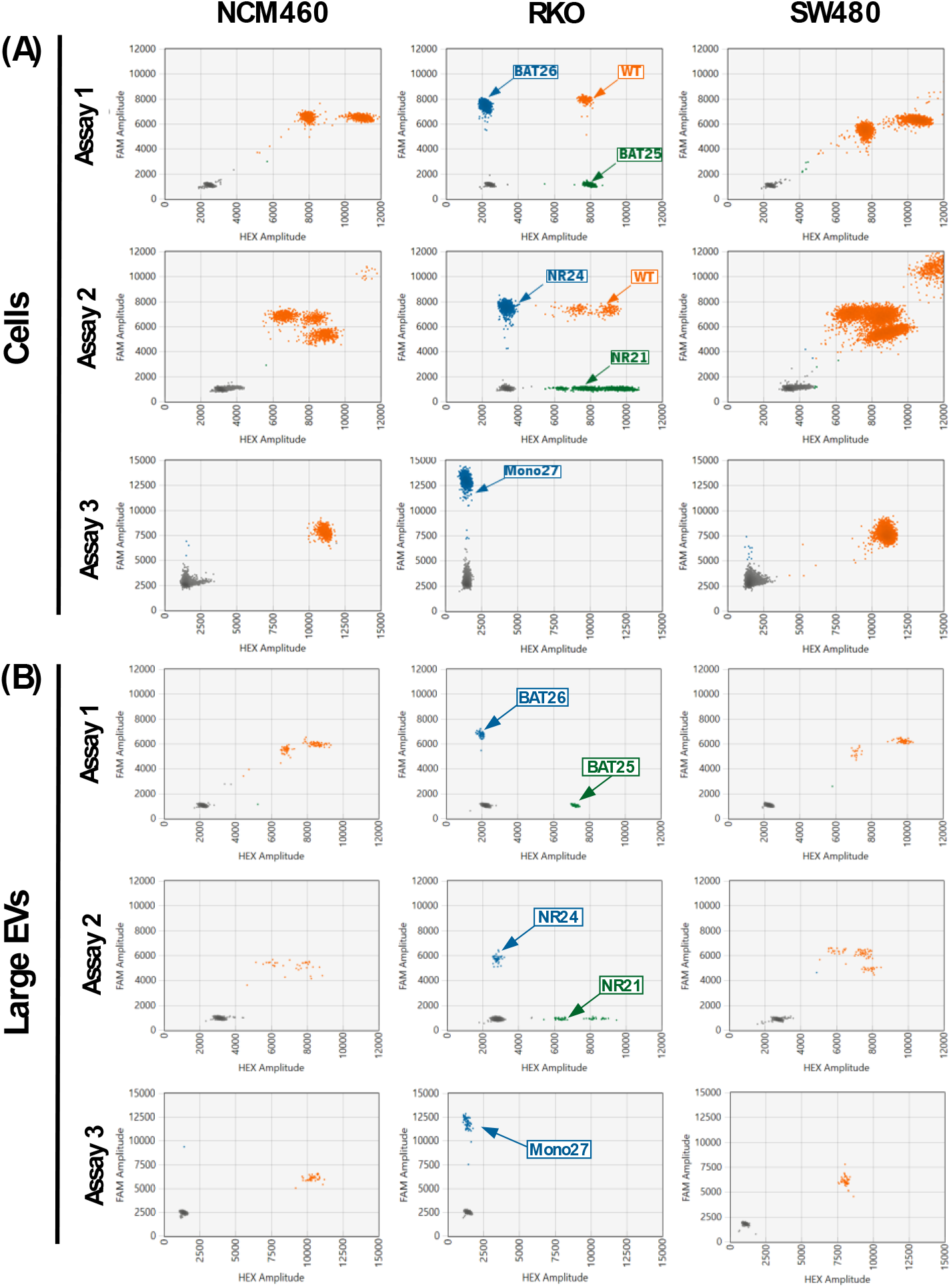
MSI detection with all assays. Detection of MSI or MSS for (A) cells, and (B) large EVs. Data analysis was performed using QX Manager Software.

Despite the documented detection of mutations (e.g. KRAS) in EV-DNA (8), evidence of MSI markers in EV-DNA, analogous to those in parent cells, has not previously been documented. This study represents the first successful detection of MSI in both L-EVs and s-EVs, even after treatment with SCFAs and with DNase application, highlighting the promise of these vesicles for liquid biopsy purposes. As illustrated in **Figure 5 A**, the markers BAT25 and BAT26 were detected in CRC cells and their respective EVs, along with NR21, NR24, and Mono27, summarized in **Figure 5 B**. This detection was possible even after treatment with SCFAs, indicating that they appear to have no effect on the expression of microsatellite instability. Notably, MSI was detected in samples where the amount of dsDNA was below the detection limit of the Qubit system, as shown in **Figure 3 B** and **C**. This underscores the capability of ddPCR to identify such characteristics even with minimal amounts of dsDNA.

**Figure 5.**
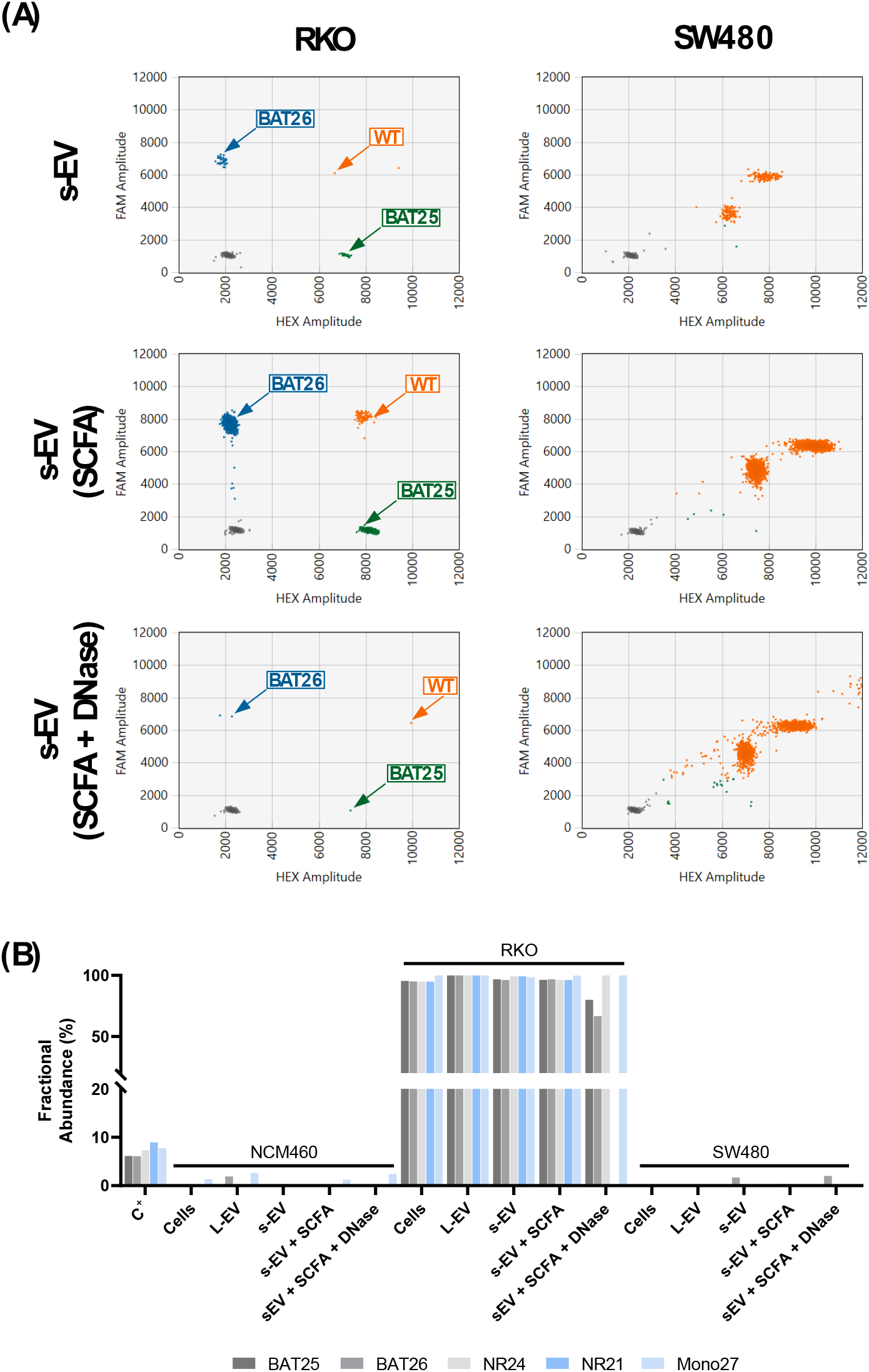
Summary of MSI detection. (A) Comparison of BAT25 and BAT26 marker detection (Assay 1) in s-EVs between an MSI CRC cell line (RKO) and a MSS CRC cell line (SW480). (B) Representation of the fractional abundances of the five markers used in the Bio-Rad ddPCR kit for MSI detection, for the three cell lines under study (NCM460, RKO, and SW480) and their respective EVs. Data analysis was performed using QX Manager Software.

These findings support EV-DNA as a viable source for MSI detection, with significant implications for liquid biopsy applications in CRC. The preservation of MSI markers within EVs, independent of DNA fragmentation and cell-free DNA limitations, enhances the potential for non-invasive diagnostics. Furthermore, the ability to detect MSI in minimal DNA quantities underscores the robustness of ddPCR and suggests a promising path forward for EV-based biomarker discovery, paving the way for EVs to become essential assets in CRC diagnostics and monitoring of tumor dynamics.

## Conclusion

This study provided new understandings into the effects of SCFAs on EV production and phenotype in CRC cells. SCFA treatment not only significantly increased the number of EVs but also influenced their nucleic acid content, particularly enhancing DNA packaging into these vesicles. These results point to the potential role of SCFA in modulating EV release during cellular responses, including stress and apoptosis.

A key finding was the successful detection of MSI markers within EV-DNA using ddPCR. This was achieved even in samples where traditional quantification methods, such as Qubit, lacked sensitivity, underscoring the potential of EVs as carriers of tumor-specific markers for liquid biopsy applications.

In conclusion, this study highlights EVs as a valuable platform for advancing non-invasive biomarker discovery in CRC and demonstrates the potential of SCFA-induced EV modifications to reflect tumor-specific characteristics. Together, these findings lay the groundwork for further exploration of EVs as diagnostic tools and for developing improved liquid biopsy methods that could transform patient management and personalized care in CRC and, potentially, other types of cancer.

## Appendix 1

**Figure A1.**
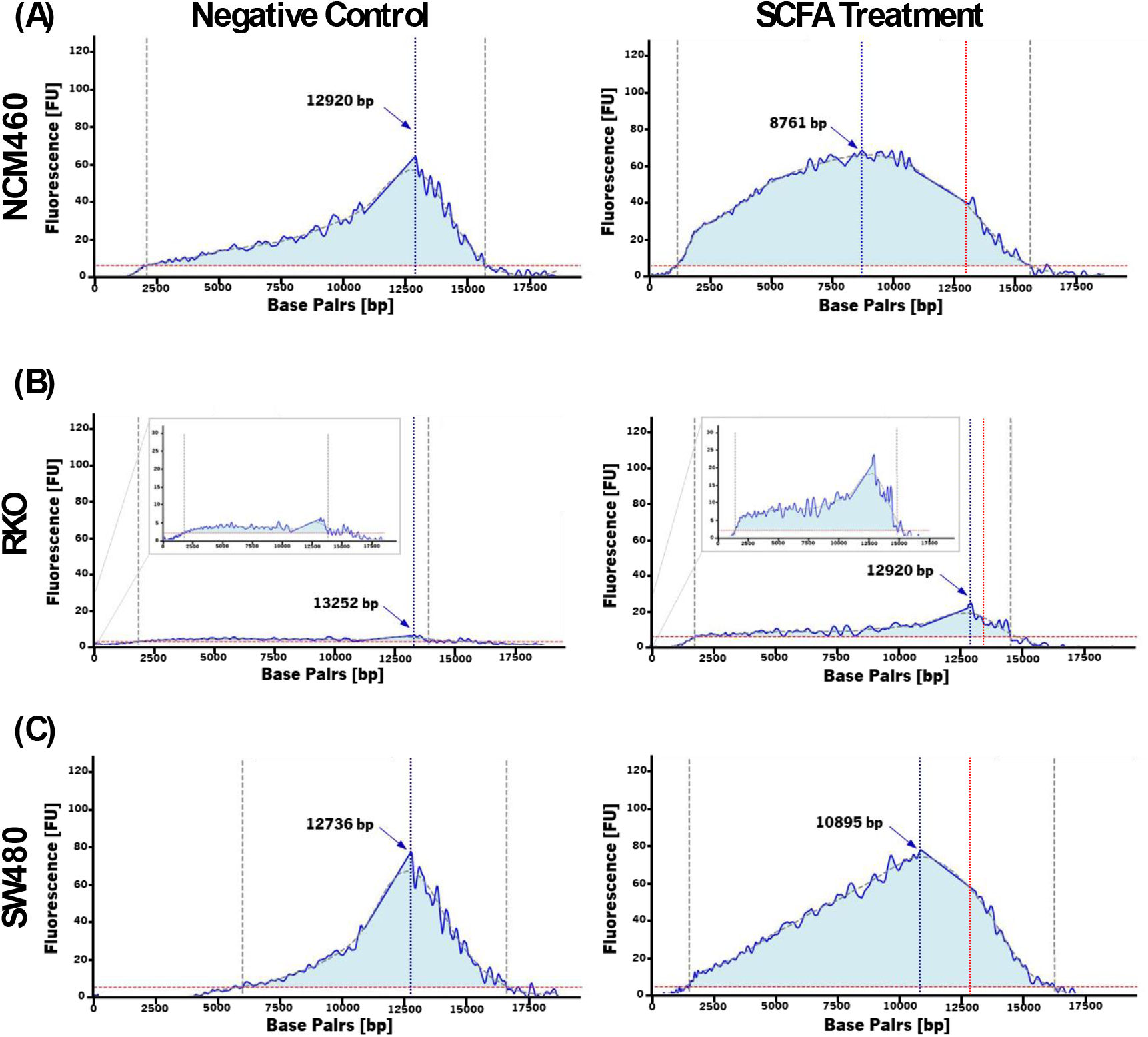
Effect of SCFA treatment on dsDNA integrity. Fragmentation profiles of dsDNA from (A) NCM460, (B) RKO, and (C) SW480 cell lines, before (left) and after treatment with SCFA (right). All samples were treated with DNase I before DNA extraction. Integrity analysis was performed using Bioanalyzer DNA 12,000 Kit and the data was further processed employing a custom Python script.

**Table A1.** Effect of SCFA treatment on dsDNA fragmentation in cells.

|  | Size range (bp) * |  | Area (%) ** |  |
| --- | --- | --- | --- | --- |
|  | Negative Control | SCFA Treatment | Negative Control | SCFA Treatment |
| **NCM460** | 2,088 – 15,719 | 1,213 – 15,608 | 70.63 | 92.89 |
| **RKO** | 1,798 – 13,840 | 1,820 – 14,504 | 90.55 | 87.79 |
| **SW480** | 6,000 – 16,545 | 1,640 – 16,271 | 55.73 | 79.30 |

\* Interval of fragmentation (in base pairs)
\*\* Percentage of the more fragmented region (left) relative to the control peak of base pairs

